# Differential serotonergic modulation of principal neurons and interneurons in the anterior piriform cortex

**DOI:** 10.1101/2022.01.05.475043

**Authors:** Ildikó Piszár, Magor L. Lőrincz

**Affiliations:** Department of Physiology, Anatomy and Neuroscience, Faculty of Sciences University of Szeged, 6726 Szeged, Hungary; Department of Physiology, University of Szeged, Szeged 6720, Hungary MTA-SZTE; ‘Momentum’ Oscillatory Neuronal Networks Research Group, Department of Physiology, University of Szeged, Szeged 6720, Hungary; Neuroscience Division, Cardiff University, Museum Avenue, Cardiff CF10 3AX, UK

## Abstract

Originating from the brainstem raphe nuclei, serotonin is an important neuromodulator involved in a variety of physiological and pathological functions. Specific optogenetic stimulation of serotonergic neurons results in the divisive suppression of spontaneous, but not sensory evoked activity in the majority of neurons in the primary olfactory cortex and an increase in firing in a minority of neurons. To reveal the mechanisms involved in this dual serotonergic control of cortical activity we used a combination of in vitro electrophysiological recordings from identified neurons in the primary olfactory cortex, optogenetics and pharmacology and found that serotonin suppressed the activity of principal neurons, but excited local interneurons. The results have important implications in sensory information processing and other functions of the olfactory cortex and related brain areas.

## INTRODUCTION

Serotonin (5-hydroxytryptamine, 5-HT) originates from neurons located in the brainstem raphe nuclei projecting to various forebrain structures including the primary olfactory cortex, the anterior piriform cortex (aPC) and is involved in various physiological and pathological phenomena including sensory and motor responses as well as the regulation of mood and impulsivity (Okaty et al., 2019). The anterior piriform cortex (aPC) receives afferent (feed-forward) inputs from the olfactory bulb (OB) and associative (feedback) inputs originating from local recurrent collaterals of aPC neurons, orbitofrontal cortex, amygdala, and entorhinal cortex (Haberly and Price, 1978; Insausti et al., 1987; Carmichael et al., 1994; Johnson et al., 2000).

5-HT can depolarize interneurons, hyperpolarize or depolarize and induce IPSPs in pyramidal neurons of the aPC (Sheldon and Aghajanian, 1990; Gellman and Aghajanian, 1994; Marek and Aghajanian, 1994, 1996) and reduces the excitability of pyramidal neurons directly (Wang et al., 2020). Importantly, specific stimulation of 5-HT neurons has a multiplicative and frequency-dependent effect on the baseline activity of aPC neurons, but no effect on odor evoked responses (Lottem et al., 2016). While the activity of most affected neurons was suppressed, a minority of neurons showed an increase in spontaneous activity following optogenetic activation of 5-HT neurons suggesting a cell-type specific action. Accordingly, in the hippocampus, the effects of 5-HT are both cell-type specific affecting cholecystokinin-expressing and O-LM interneurons, but not parvalbumin-expressing basket cells, and input specific, affecting only glutamatergic synaptic transmission originating from CA1 pyramidal cells (Winterer et al., 2011; Bohm et al., 2015).

As the cell-type specific effects are difficult to reveal from the extracellular action potentials recorded *in vivo* we sought to identify these effects using a combination of electrophysiological, pharmacological and optogenetic methods in vitro. We found that aPC interneurons including perisomatic inhibitory fast spiking interneurons are excited by 5-HT whereas principal neurons are inhibited. These results could have important implications for aPC function.

## MATERIALS AND METHODS

A cre-dependent adeno-associated viral vector AAV1-ChR2-YFP (AV-1-20298P, University of Pennsylvania, 10^13^ GC/mL) was injected into the DRN of adult BAC transgenic Slc6a4-cre (i.e. SERT-cre) mice (Gong et al., 2007) as previously described (Dugue et al., 2014). 8-15 weeks following the viral infections mice were used for *in vitro* experiments. All animals used in this study showed prominent somatic YFP fluorescence in the DRN region and axonal YFP fluorescence in the aPC (Figure 2B). For focal 5-HT application experiments wild type littermates of SERT-cre mice were used. Brain slices were prepared similar to previously described protocols (Gazea et al., 2021). Briefly, mice were deeply anesthetized with ketamine and xylazine (80 and 10 mg/kg, respectively), and perfused through the heart with a solution containing (in mM): 93 NMDG, 2.5 KCl, 1.2 NaH_2_PO_4_, 30 NaHCO_3_, 20 HEPES, 25 glucose, 5 N-acetyl-L-cysteine, 5 Na-ascorbate, 3 Na-pyruvate, 10 MgSO_4_, and 0.5 CaCl_2_. The same solution was used to cut 320 μm coronal slices containing the aPC at 4°C and for the initial storage of slices (32°C-34°C for 12 min) following which the slices were stored in a solution containing the following (in mM): 30 NaCl, 2.5 KCl, 1.2 NaH_2_PO_4_, 1.3 NaHCO_3_, 20 HEPES, 25 glucose, 5 N-acetyl-L-cysteine, 5 Na-ascorbate, 3 Na-pyruvate, 3 CaCl_2_, and 1.5 MgSO_4_. For recording, slices were submerged in a chamber perfused with a warmed (34°C) continuously oxygenated (95% O2, 5%CO2) ACSF containing the following (in mM): 130 NaCl, 3.5 KCl, 1 KH_2_PO_4_, 24 NaHCO_3_, 1.5 MgSO_4_, 3 CaCl_2_, and 10 glucose. Whole-cell patch clamp recordings were performed in the current-clamp mode using 4-6MOhm pipettes containing (in mM): 126 Kgluconate, 4 KCl, 4 ATP-Mg, 0.3 GTP-Na2, 10 HEPES, 10 creatine-phosphate, and 8 Biocytin, pH 7.25; osmolarity, 275 mOsm. Neurons were visualized using DIC imaging on an Olympus BX51WI microscope (Tokyo, Japan). Membrane potentials were recorded using a Multiclamp 700B amplifier (Molecular Devices, USA). The liquid junction potential (−13 mV) was compensated for. Series resistance was continuously monitored and compensated during the course of the experiments; recordings were discarded if the series resistance changed more than 25%. For the focal application of 5-HT patch pipettes were loaded with 100 μM 5-HT dissolved in ACSF, positioned near (40-60 μm) the soma of the neuron recorded and connected to a Picospritzer III (Parker Hannifin, USA). 5-HT was ejected using a 2 s long (~200 mbar) pulse. The recorded neurons were classified as pyramidal neurons based on somatic morphology under DIC (presence of a prominent apical dendrite, pyramidal shaped somata in layer 2 or 3), membrane responses to hyperpolarizing and depolarizing current steps including a relatively low input resistance or axodendritic arborizations following post-hoc fluorescent imaging following Streptavidin immunoreactions (Iacone et al., 2021). The recorded neurons were classified as interneurons based on somatic morphology under DIC (absence of a prominent apical dendrite, small, round or oval somata in layers 1-3), membrane responses to hyperpolarizing and depolarizing current steps including a relatively high input resistance and presence of EPSPs or axodendritic arborizations following post-hoc fluorescent imaging following Streptavidin immunoreactions. Immunostaining of 5-HT fibers in the aPC was performed as previously described (Varga et al., 2009). Briefly, 320 μm coronal brain slices used for in vitro recordings were fixed in 4% paraformaldehyde overnight at 4 °C. The slices were resectioned to 60 μm and incubated in 5% NGS and 0.1% Triton and the primary antibody (Immunostar, Hudson, WI, USA, Rb-α-5-HT polyclonal, 1:1000). Alexa488-conjugated Donkey-α-Mouse (1:200) was used as a secondary antibody and the slices mounted in VectaShiled medium for microscopy. Fluorescent images were acquired with a confocal microscope (Olympus FV1000). Data are presented as mean±s.e.m. All drugs were purchased from Tocris Biosciences (Bristol, UK). Concentration of the drugs used: ketanserin: 10 μM, WAY100635 1 μM.

## RESULTS

To characterize the extent of serotonergic innervation of the aPC we performed immunhistochemical experiments by staining the 5-HT fibers with an antibody against 5-HT in coronal brain slices containing the aPC (see Materials and Methods). The results revealed dense 5-HT fibers in the aPC with subregion and layer specific features (Figure 1A). Specifically, the aPC contained relatively dense 5-HT fibers in all its layers, most fibers were observed in layer 1 and 3. To test the effect of 5-HT on various aPC neurons we focally applied 5-HT while monitoring their membrane potential (Figure 1A). Morphologically and physiologically identified pyramidal neurons were hyperpolarized by 5-HT (control: - 70.47±0.79, 5-HT: −73.38±1.07, p<0.05, Wilcoxon signed-rank test, n=5) (Figure 1B, C and D). When the membrane potential of neurons was held around the threshold for action potential generation by injecting steady depolarizing current via the recording electrode, the application of 5-HT suppressed their firing (Figure 1B). Both the hyperpolarization (5-HT: - 2.64±0.47 mV, WAY100635 + 5-HT: 0.05±0.14 mV, p<0.05, Wilcoxon signed-rank test, n=5) and the suppression of action potential firing could be prevented by the bath application of the 5-HT1a receptor antagonist WAY 100635 (5-HT: 1.58±1.01%, WAY100635 + 5-HT: 108.93±1.57%, p<0.001, Wilcoxon signed-rank test n=5) suggesting the effect of 5-HT on pyramidal neurons is mediated by 5-HT1 receptors (Figure 1B, C and D). When 5-HT was focally applied near the somata of various identified interneurons while monitoring their membrane potential, the cells were depolarized (5-HT: 7.30±3.24 mV, n=5) and this depolarization led to action potential firing in all the recorded interneurons (mean firing rate: 2.66±1.40 Hz, n=5) (Figure 1E). Both the membrane depolarization and the action potential firing could be prevented by the bath application of the 5-HT2 receptor antagonist ketanserin (Figure 1E and F) suggesting the effect of 5-HT on interneurons is mediated by 5-HT2 receptors.

**Figure 1.**
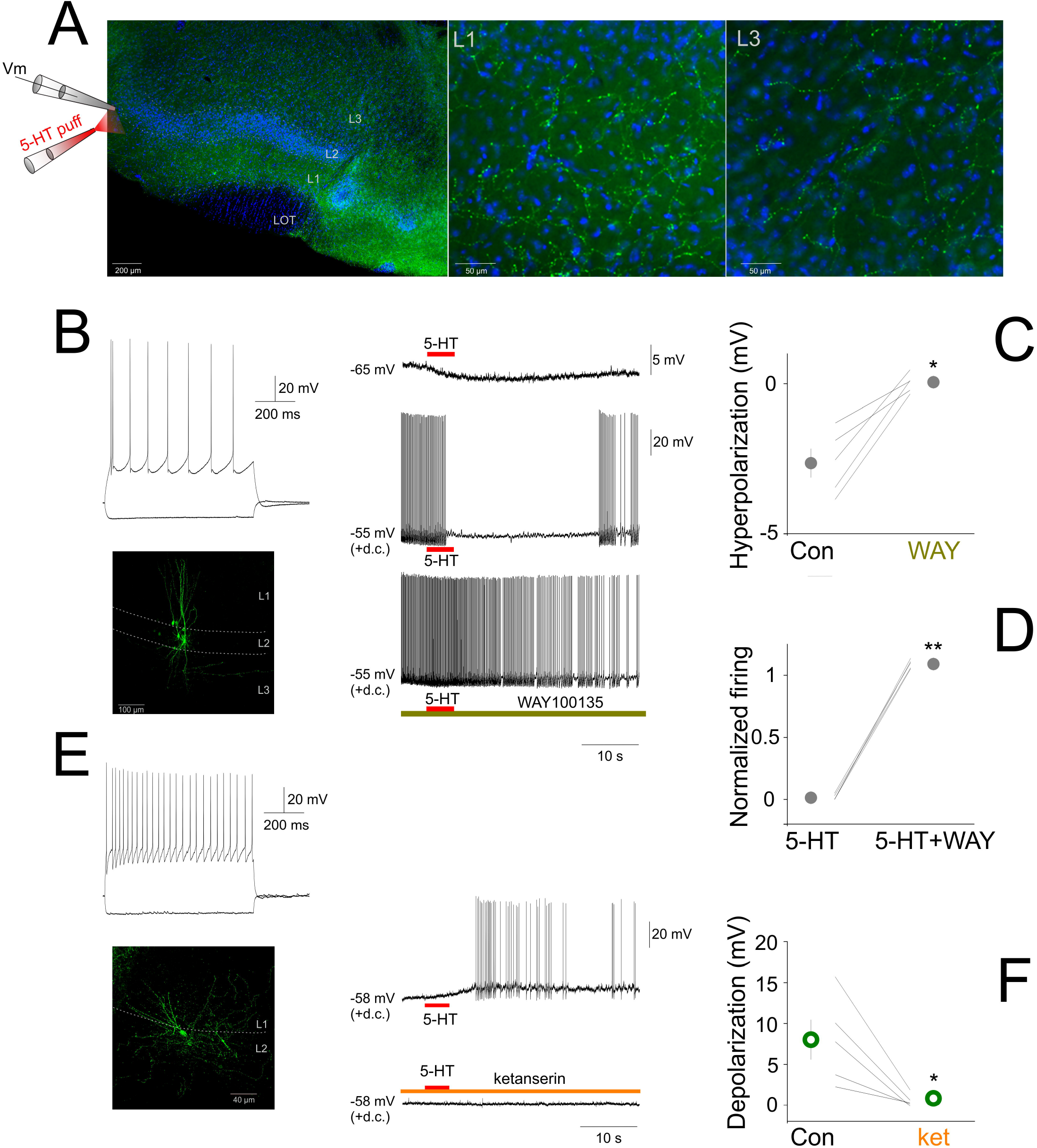
Focal 5-HT application suppresses the activity of principal neurons and increases the activity of interneurons in the aPC. (A) Schematic of the experimental design and serotonergic axonal innervation (green channel: serotonin immunopositive fibers) of the aPC. Coronal section of a mouse brain at the level of the aPC. The lateral olfactory tract (LOT) and the three layers of the aPC are indicated. Blue channel: DAPI staining. Higher magnification images of layers 1 and 3, respectively are shown on the right side. (B) Membrane responses to hyperpolarizing and depolarizing current steps (top left), morphology (bottom left) and effect of focally applied 5-HT (100 μM, middle) of a principal neuron at resting membrane potential and during periods of action potential firing (+d.c.). The 5-HT1a antagonist WAY100635 (1 μM) prevented the suppressive effects of 5-HT. (C) Effects of focally applied 5-HT in ACSF (Con) and WAY 100635 (WAY) on all recorded aPC principal neurons (n=5). (D) Normalized firing rates of all recorded principal neurons (n=5) during focal 5-HT application in the absence and presence of WAY 100635 (WAY), respectively. (E) Membrane responses to hyperpolarizing and depolarizing current steps (top left), morphology (bottom left) and effect of focally applied 5-HT (100 μM, middle) of an aPC interneuron. The 5-HT2 antagonist ketanserin (10 μM) prevented the depolarizing effects of 5-HT. (F) Effects of focally applied 5-HT in ACSF (Con) and ketanserin (ket) on all recorded aPC interneurons (n=5).

Since the effects of exogenously applied 5-HT can differ from the effects of endogenous 5-HT, we next tested the effects of endogenously released 5-HT on various aPC neurons by monitoring their membrane potential and selectively stimulating local 5-HT axons in SERT-cre mice previously (8-15 weeks) infected with ChR2 in the DRN (Dugue et al., 2014), which led ChR2 expression in the somata of DRN 5-HT neurons and prominent axonal ChR2-YFP expression in the aPC (Figure 2B). In 20 morphologically and/or physiologically identified pyramidal neurons the local photostimulation (PS) of 5-HT axons did not lead to an apparent membrane potential hyperpolarization (mean Vm change following 5-HT PS: −0.03±0.04 mV, p>0.05, Wilcoxon signed-rank test, n=20) (Figure 2C). When we photostimulated 5-HT axons while monitoring the membrane potential of various morphologically and/or physiologically identified interneurons we observed a membrane potential depolarization in 5 out of 8 (62%) interneurons (mean Vm change following 5-HT PS: 1.11±0.42 mV, n=8). 5-HT axonal PS in the aPC could lead to action potential firing in 4 out of 8 (50%) interneurons recorded (Figure 2E). Both the depolarization and the effect on action potential firing could be prevented by bath application of the 5-HT2 receptor antagonist ketanserin (10 μM) (Figure 2E) suggesting aPC interneurons can be excited by endogenously released 5-HT originating from the axons of DRN 5-HT neurons via 5-HT2 receptors.

**Figure 2.**
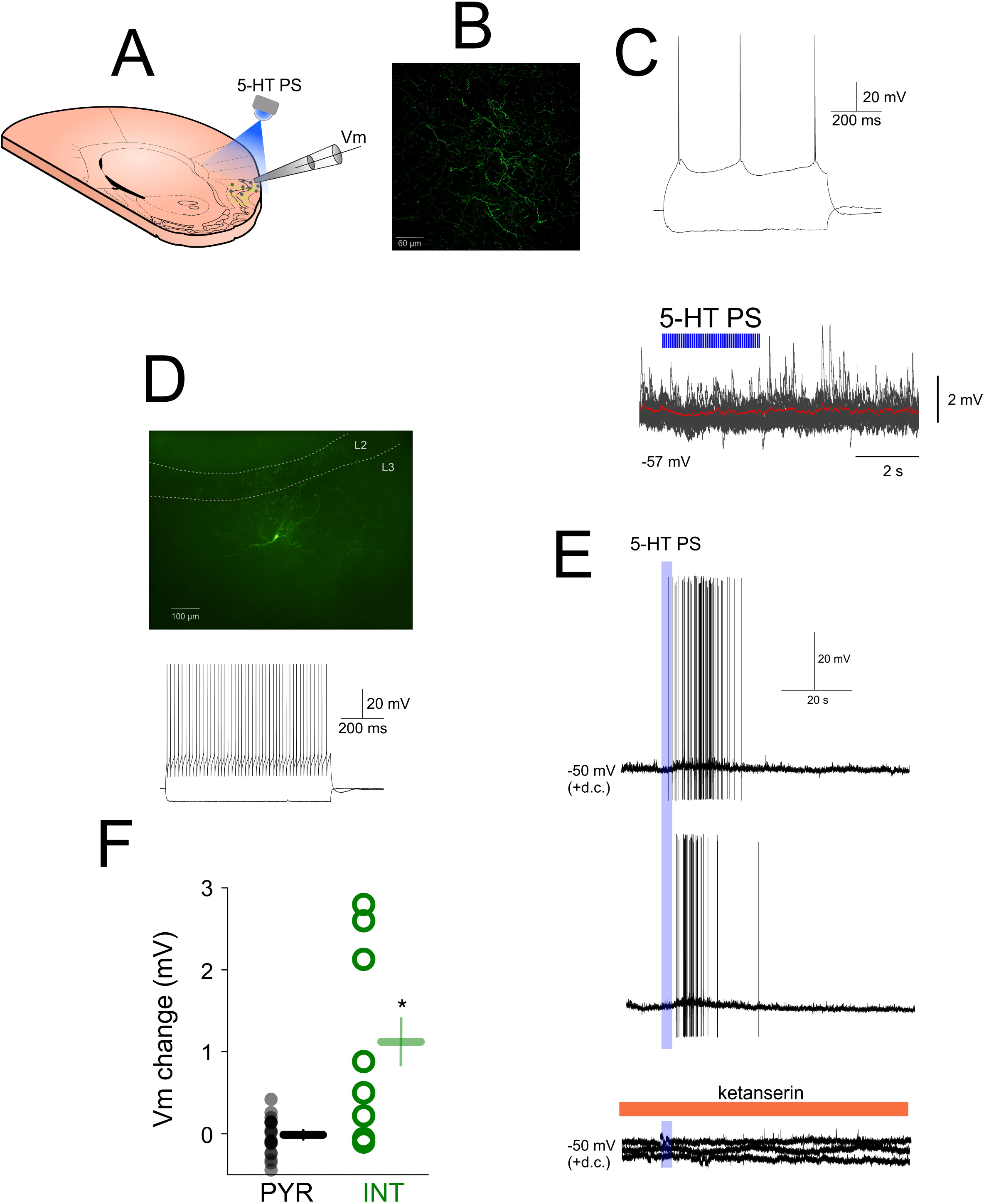
Effects of endogenously released 5-HT on aPC principal neurons and interneurons. (A) Schematic of the experimental design. (B) Confocal image of ChR2-YFP expressing fibers in the aPC of a SERT-cre mouse previously injected with AAV-DIO-ChR2-eYFP in the DRN. (C) Membrane potential responses of an aPC pyramidal neuron to hyperpolarizing and depolarizing current steps (top) and lack of effect of local photostimulation (10 ms pulses, 20 Hz, 5 mW, 3 s) of DRN 5-HT axons in the aPC on the membrane potential (20 sweeps overlayed, red trace: average) of the aPC pyramidal neuron shown in B. (D) Morphology (top) of an aPC fast spiking interneuron and its membrane responses to hyperpolarizing and depolarizing current steps (bottom). (E) Effect of local photostimulation (10 ms pulses, 5 mW, 20 Hz, 3 s) of DRN 5-HT axons in the aPC on the membrane potential of the neuron shown in C (two consecutive sweeps). The depolarizing effect is blocked by ketanserin (three consecutive sweeps). (F) Effects of 5-HT photostimulation on all recorded aPC pyramidal neurons (PYR, gray circles, n=20) and interneurons (INT, green circles, n=7).

The presence of a direct membrane effect on interneurons and the lack of effect on pyramidal neurons, respectively of the endogenously released 5-HT suggested that 5-HT could control the rate and timing of action potential output of aPC pyramidal neurons and interneurons as previously reported (Wang et al., 2020). To test this, we applied suprathreshold depolarizing current steps (50-200 pA) and compared the mean firing rates evoked by the same current pulse in various aPC neurons in the absence and the presence of 5-HT PS, respectively (Figure 3). We found that the evoked firing rates were significantly lower in pyramidal neurons in the presence of 5-HT PS (normalized mean firing rate during 5-HT PS: 30.96±9.51%, p<0.01, Wilcoxon signed-rank test, n=16) (Figure 3A and D). On the other hand, when we repeated the same protocol in aPC interneurons we found that the evoked firing rates of 7 interneurons out of 10 (70%) were higher in the presence of 5-HT phostimulation, although this was not significant for the whole group (normalized mean firing rate during 5-HT PS: 133.73±28.63%, p>0.05, Wilcoxon signed-rank test, n=10, Figure 3B, C and D). These results suggest that the endogenously released 5-HT can affect the activity of aPC neurons in a cell type specific manner, boosting the activity of interneurons and suppressing pyramidal neuron firing.

**Figure 3.**
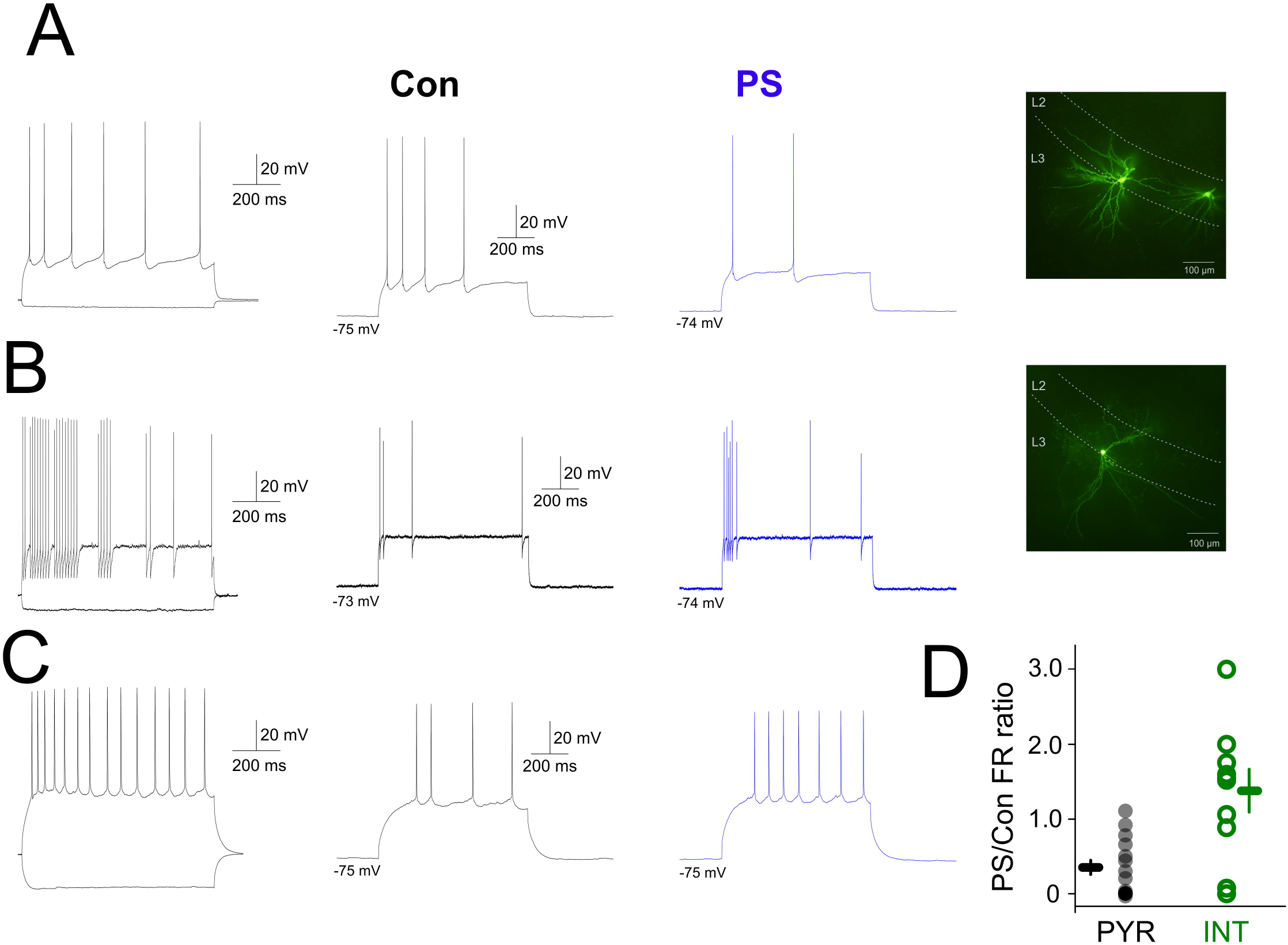
Effects of endogenously released 5-HT on the evoked firing of aPC principal neurons and interneurons. (A) Membrane potential responses and morphology of an aPC pyramidal neuron to hyperpolarizing and depolarizing current steps in the absence (Con, black trace) and presence (PS, blue trace) of 5-HT photostimulation, respectively. (B) Membrane potential responses and morphology of an aPC interneuron to hyperpolarizing and depolarizing current steps in the absence (Con, black trace) and presence (PS, blue trace) of 5-HT photostimulation, respectively. (C) Same as D for a regular firing aPC interneuron. (D) Firing (normalized to control) of all photostimulated pyramidal neurons (PYR, gray circles, n=20) and interneurons (INT, green circles, n=8).

## DISCUSSION

Using a combination of in vitro electrophysiology, optogenetics and pharmacology we found that aPC interneurons including perisomatic inhibitory fast spiking interneurons are excited and principal neurons inhibited by 5-HT. The results of 5-HT focal application show a prominent hyperpolarization of aPC principal neurons in accordance with previous studies using bath applied 5-HT (Sheldon and Aghajanian, 1990; Gellman and Aghajanian, 1994; Marek and Aghajanian, 1994, 1996). Interestingly, endogenously released 5-HT from ChR2 expressing DRN 5-HT axons had no prominent hyperpolarizing effect on the membrane potential of aPC principal neurons. While this could be caused by ineffective release of 5-HT from DRN axons in the aPC an alternative explanation is that the exogenously applied 5-HT can activate receptors that are far from any 5-HT terminal as in the case of dopamine (Rosen et al., 2015). In line with this, endogenous 5-HT release did affect the membrane potential of some aPC interneurons and the evoked firing of both principal neurons and interneurons. Taken together these results argue against the possibility of a failure in 5-HT release from ChR2 expressing DRN terminals in vitro and suggest that 5-HT has a subtle cell-type specific effect on aPC neurons.

These results are in line with our previous observations showing a suppressive effect of 5-HT photostimulation on the spontaneous activity of most aPC neurons and an increase in firing in a minority of aPC neurons (Lottem et al., 2016). In addition to the direct suppressive effects of 5-HT on principal neurons, the increase in activity in various interneurons, including perisomatic targeting fast spiking interneurons could be an additional mechanism by which 5-HT can potently control the activity of aPC principal neurons. Whether the same DRN neurons target multiple types of aPC neurons including principal neurons and interneurons or whether the connectivity itself possesses cell-type specific features remains to be established. As the activity of DRN neurons is modulated on relatively rapid timescales in a behaviorally relevant manner (Lőrincz et al., 2007; Ranade and Mainen, 2009; Liu et al., 2014; Cohen et al., 2015; Fonseca et al., 2015; Lottem et al., 2018) it is interesting to speculate on the potential consequences of rapid and cell-type specific 5-HT effects on aPC neurons. In addition to potential effects on central and peripheral sensory information processing (Lőrincz et al., 2008; Kapoor et al., 2016; Lottem et al., 2016) 5-HT could also control the timing of various aPC neurons in relation to gamma oscillations originating in the olfactory bulb which are known to be important in widespread gamma oscillations in the limbic system (Becker and Freeman, 1968). Interestingly, fast spiking interneurons, the more strongly affected neuronal population in our 5-HT PS experiments are key players in the generation and maintenance of cortical gamma oscillations (Cardin et al., 2009). As limbic gamma oscillations are important in the maintenance of a healthy mood and are affected in major depression (Scangos et al., 2021) the effects of 5-HT described here could be also relevant for the (patho)physiology of higher brain functions.

## Author contributions

MLL and IP designed the experiments, IP performed the experiments, IP and MLL analysed the data and prepared the figures. MLL and IP wrote the manuscript. All authors contributed to the article and approved the submitted version.

## Funding

This work was supported by the Hungarian Scientific Research Fund (Grants NN125601 and FK123831 to M.L.L.), the Hungarian Brain Research Program (grant KTIA_NAP_13-2-2014-0014, UNKP-20-5 New National Excellence Program of the Ministry for Innovation and Technology from the source of the National Research, Development and Innovation Fund to MLL. MLL is a grantee of the János Bolyai Fellowship.

## Conflict of interest

The authors declare no conflict of interest.

## Notes

### Competing Interest Statement

The authors have declared no competing interest.

